# The activity of human enhancers is modulated by the splicing of their associated lncRNAs

**DOI:** 10.1101/2020.04.17.045971

**Authors:** Jennifer Y. Tan, Ana C. Marques

## Abstract

Pervasive enhancer transcription is at the origin of more than half of all long noncoding RNAs in humans. Transcription of enhancer-associated long noncoding RNAs (elncRNA) contribute to their cognate enhancer activity and gene expression regulation in *cis.* Recently, splicing of elncRNAs was shown to be associated with elevated enhancer activity. However, whether splicing of elncRNA transcripts is a mere consequence of accessibility at highly active enhancers or if elncRNA splicing directly impacts enhancer function, remains unanswered.

We analysed genetically driven changes in elncRNA expression, in humans, to address this outstanding question. We showed that splicing related motifs within multi-exonic elncRNAs evolved under selective constraints during human evolution, suggesting the processing of these transcripts is unlikely to have resulted from transcription across spurious splice sites. Using a genome-wide and unbiased approach, we used nucleotide variants as independent genetic factors to directly assess the causal relationship that underpin elncRNA splicing and their cognate enhancer activity. We found that the splicing of most elncRNAs is associated with changes in chromatin signatures at cognate enhancers and target mRNA expression.

We conclude that efficient and conserved processing of enhancer-associated elncRNAs contributes to enhancer activity.

## INTRODUCTION

Classically, enhancers are defined as regulatory DNA elements that positively regulate temporal and spatial expression of their target genes, in *cis*. Transcription activation by enhancers requires transcription factor-dependent recruitment of coactivating complexes and three-dimensional genome rearrangements that bring enhancers into close proximity of their target promoters [reviewed in (Schoenfelder and Fraser 2019)].

Whereas enhancer activity, as classically defined, is solely determined by its sequence and genomic location, most, if not all, active enhancers are transcribed (De Santa et al. 2010; Kim et al. 2010; Kowalczyk et al. 2012). Initial evidence of this phenomenon emerged from the characterisation of some of the first identified enhancers (Ashe et al. 1997). Pervasive transcription of active enhancers was later confirmed genome-wide, initially in in neurons (Kim et al. 2010) and macrophages (De Santa et al. 2010) and more recently across a number of human and mouse cell types (Andersson et al. 2014; Arner et al. 2015). These studies revealed that enhancer transcription often precedes target promoter activation (De Santa et al. 2010) and is positively correlated with target gene expression (De Santa et al. 2010; Kim et al. 2010; Arner et al. 2015). The majority [>95% (Natoli and Andrau 2012; Gil and Ulitsky 2018; Tan et al. 2020)] of enhancers is transcribed bidirectionally and gives rise to relatively short single exonic RNAs that are non-polyadenylated and unstable, generally referred to as eRNAs (Kim et al. 2010). The remaining enhancers transcribe enhancer-associated long noncoding RNAs [elncRNAs (Marques et al. 2013; Gil and Ulitsky 2018; Tan et al. 2020)] predominantly in one direction (Natoli and Andrau 2012). In contrast with eRNAs, elncRNAs are relatively stable, polyadenylated and can be spliced (Natoli and Andrau 2012).

Transcription has been found to strengthen cognate enhancer activity through various non-mutually exclusive mechanisms. For example, enhancer transcription facilitates binding of enhancer factors, such as CREBBP (Bose et al. 2017), or chromatin remodeling complex, including Cohesin and Mediator, that in turn induce RNA-dependent changes in local chromatin conformation (Lai et al. 2013; Mousavi et al. 2013; Hsieh et al. 2014). Enhancer transcription has also been proposed to regulate the load, pause and release of RNA Polymerase II (RNAPlI) (Maruyama et al. 2014; Schaukowitch et al. 2014).

While the contribution of transcription to enhancer function is by now relatively well established, the importance of the transcribed RNAs has only been demonstrated for a handful of anecdotal transcripts (for example, (Lai et al. 2013; Lam et al. 2013; Li et al. 2013; Melo et al. 2013; Iott et al. 2014)). In general, the absence of evolutionary constraint at eRNA/elncRNA exons (Marques et al. 2013) argues that the function of most eRNAs/elncRNAs is unlikely to be encoded within their sequences. Yet a large fraction of elncRNAs is multi-exonic and recently, genome-wide analysis in multiple human cell lines (Gil and Ulitsky 2018) and mouse Embryonic Stem Cells (Tan et al. 2020) revealed that their splicing positively impacts enhancer function. Enhancers that transcribe multi-exonic elncRNAs are enriched in enhancer-specific chromatin signatures, elevated binding of co-transcriptional regulators, increased local intra-chromosomal DNA contacts, and strengthened *cis*-regulation on target gene expression (Gil and Ulitsky 2018; Tan et al. 2020). What remains currently unclear is the causal relationship between elncRNA splicing and enhancer activity. High enhancer activity may promote transcription across splice/donor acceptor site-like sequence, which are common in mammals (Wan and Larson 2018), leading to unregulated/noisy splicing of elncRNAs. Arguing against this is evidence that mutations disrupting elncRNA splicing directly decreased their cognate enhancer activity. For example, removal of the intronic sequence of the locus encoding for the Haunt elncRNA significantly reduced its *cis* regulatory function (Yin et al. 2015), whereas mutations disrupting the splicing of Blustr, another enhancer-associated lncRNA, were sufficient to negatively impact the expression of its *cis* target (Engreitz et al. 2016). Whether the splicing of most elncRNAs directly impacts enhancer activity, as exemplified by Blustr and Haunt, or whether it is an inconsequential result of their cognate enhancer’s high activity remains unknown.

We used a combination of integrative and population genomics approaches to address this outstanding question. We showed that splicing related motifs within elncRNAs evolved under selective constraint, consistent with their processing being biologically relevant. Using a mediation-based approach, we performed causal inference testing using genetic variants as independent factors to test the underlying causal relationship between elncRNA splicing and their cognate enhancer function. We showed that splicing of most elncRNAs causally contribute to target gene regulation. Furthermore, nucleotide variants that disrupt elncRNA splicing significantly reduced cognate enhancer activity, supporting the functional importance of elncRNA splicing.

## RESULTS

### Splicing of elncRNAs is associated with higher enhancer activity

We integrated publicly available RNA sequencing data for human lymphoblastoid cell lines [LCL (Lappalainen et al. 2013)] with enhancer annotations from the ENCODE consortium (Hoffman et al. 2013) to distinguish between predominantly bidirectionally (eRNAs, n=4433) and unidirectionally transcribed (elncRNAs, n=564) enhancers active in LCLs (Supplementary Table ST1, Methods). The transcription profiles of elncRNAs resemble that of promoter-associated lncRNAs (plncRNAs, n=600) and protein-coding genes (n=12,070, Supplementary Figure S1A). In LCLs, most (n=352, 62.4%) elncRNAs are multi-exonic, consistent with previous analysis in other human cell lines (Gil and Ulitsky 2018).

As described for other human and mouse cells, enhancers that transcribe multi-exonic elncRNAs are more active (Gil and Ulitsky 2018; Tan et al. 2020). In particular, compared to enhancers that give rise to either single-exonic elncRNAs or eRNAs, those transcribing multi-exonic elncRNAs have higher levels of chromatin modifications associated with enhancer function (Figure 1A-C), including mono-methylation of Histone 3 Lysine 4 (H3K4me1, Figure 1A), acetylation of Histone 3 lysine 27 (H3K27ac, Figure 1B) and DNase I accessibility (DHSI, Figure 1C). Multi-exonic elncRNA-transcribing enhancers are also enriched for binding of histone acetyl transferase (HAT) P300 (Figure 1D), whose recruitment also distinctly marks active enhancers (Merika et al. 1998).

**Figure 1.**
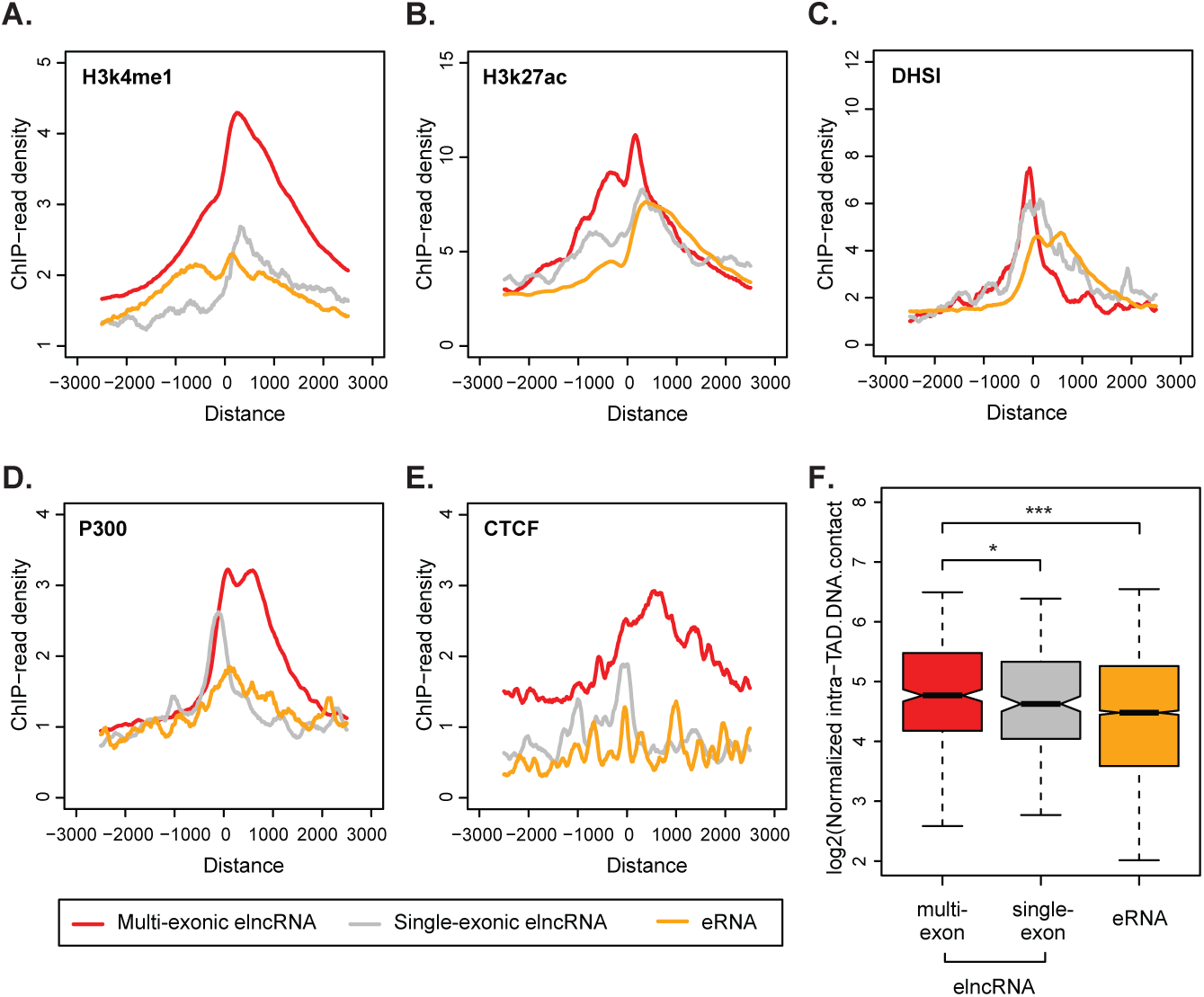
Multi-exonic elncRNAs are transcribed from highly active enhancers. Metagene plots of (A) H3K4me1, (B) H3K27ac, (C) DNase I hypersensitive sites (DHSI), (D) P300 and (E) CTCF ChIP-seq reads in LCLs at promoters of multi-exonic (red) and single-exonic (grey) elncRNAs, and eRNAs (yellow). (F) Distribution of the average amount of DNA:DNA contacts within LCL TADs containing multi-exonic (red) and single-exonic (grey) elncRNAs and eRNAs (yellow). Differences between groups were tested using a two-tailed Mann-Whitney *U* test. * *p* < 0.05; *** *p* < 0.001.

Consistent with previous analysis (Gil and Ulitsky 2018; Tan et al. 2020), relative to all other transcribed enhancers, multi-exonic elncRNA enhancers are enriched in CTFC binding (Figure 1E), significantly enriched at LCL promoter-enhancer loop domains (fold enrichment=1.3, FDR=7×10^−3^, Methods) and their topologically associating domains (TAD) display a significantly higher frequency of intra-TAD DNA-DNA contacts (p<0.02, two-tailed Mann-Whitney *U* test, Figure 1F).

In summary, our analysis in LCLs supports that enhancers transcribing multi-exonic elncRNAs are more active, as previously described in other cell types (Gil and Ulitsky 2018; Tan et al. 2020).

### elncRNA splicing was preserved by purifying selection

The splicing of elncRNAs may be a regulated and conserved process. Alternatively, it can be a mere by-product of inconsequential transcription across splice-site donor/acceptors favoured by increased accessibility at their highly active cognate enhancers. To distinguish between these two possibilities, we first analysed splicing related motifs of elncRNAs. As previously described (Schuler et al. 2014; Haerty and Ponting 2015; Tan et al. 2020) the GC contents of exons and introns (Amit et al. 2012), a determinant of efficient splice-site recognition, of multi-exonic elncRNAs is similar to that of protein-coding genes and promoter-associated plncRNAs, and as observed in mESCs (Tan et al. 2020) (Supplementary Figure S2A, two-tailed Mann-Whitney test). Compared to plncRNAs, regions flanking elncRNA splice-sites are enriched in splicing-associated elements, including U1 snRNP binding motifs (Supplementary Figure S2B, p=0.05) and a comprehensive (Supplementary Figure S2C, p=0.04) (Fairbrother et al. 2002) and a stringent set (Supplementary Figure S2D, p=0.05, two-tailed Mann-Whitney test)(Caceres and Hurst 2013) of exonic splicing enhancers (ESEs). This increase in density of splicing-related motifs in elncRNAs support their increased splicing efficiency relative to plncRNAs (Supplementary Figure S2E, p=5×10^−11^, two-tailed Mann-Whitney *U* test), consistent with what was previously reported in mESCs (Tan et al. 2020).

If splicing of elncRNAs is important for enhancer function, one would expect selection to have prevented the accumulation of deleterious mutations in their splicing-associated motifs during evolution. We estimated the nucleotide substitution rate, between human and mouse, for elncRNA splicing-related sequence motifs and compared it to that of neutrally evolving neighbouring ancestral repeats (ARs, Figure 2A). We found that splice donor/acceptor sites, as well as U1 and ESE sites, evolved under significant selective constraint by accumulating fewer substitutions during mammalian evolution than neutrally evolving ARs (p<0.002, permutation test, Figure 2A). Consistent with the observed differences in splicing efficiency, we found that splicing-associated motifs within elncRNAs (including SS, U1 and ESEs, Figure 2B-D), evolved significantly slower (p<2.2×10^−16^, two-tailed Mann-Whitney *U* test) than those within plncRNAs.

**Figure 2.**
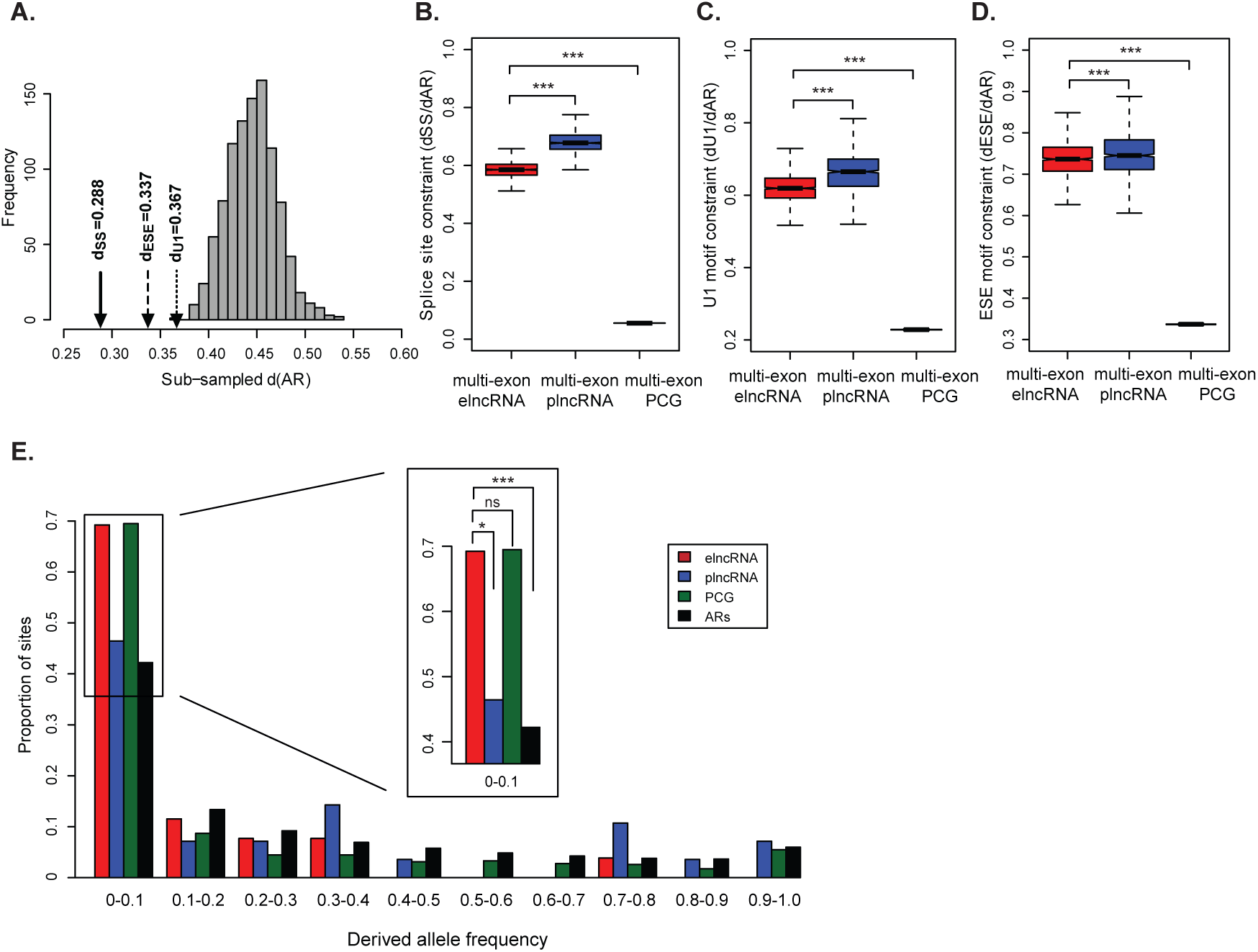
Splicing of elncRNAs has evolved under purifying selection. (A) Distribution of pairwise nucleotide substitution rate between human and mouse for 1,000 randomly subsampled sets of local ancestral repeats (ARs) with matching GC-content and size as splicing related motifs (grey). The observed rate observed at splice sites (d_SS_, solid line arrow), ESE (d_ESE_, dash line arrow), and U1 (d_U1_, dotted line arrow) sites within multi-exonic elncRNAs. Distribution of nucleotide substitution rate relative to that of randomly subsampled local ARs (d_AR_) for (B) splice sites, (C) ESEs, and (D) U1s within multi-exonic elncRNA (red), plncRNA (blue) and protein-coding gene (dark green). Differences between groups were tested using a two-tailed Mann-Whitney *U* test. *** *p* < 0.001. (E) Distribution of derived allele frequency (DAF) for single nucleotide variants at splicing motifs (splice sites, ESEs and U1s) within multi-exonic elncRNAs (red), plncRNAs (blue), protein-coding genes (dark green), and ARs (black). The insert illustrates variants with low derived allele frequency (DAF<0.1). Differences between groups were tested using a two-tailed Fisher’s exact test. * *p* < 0.05; *** *p* < 0.001; NS *p* > 0.05.

We next turned our attention to the evolution of elncRNA splicing during recent human history. Relative to neutrally evolving ARs and plncRNAs, single nucleotide polymorphisms (SNPs) within elncRNA splice sites and splicing motifs were enriched in alleles with low derived allele frequency (DAF < 0.1) (p<0.05, two-tailed Fisher’s exact test, Figure 2E), consistent with their evolution under selective constraint. Whereas, the frequency of SNPs with DAF<0.1 that overlap splicing-associated sequences within elncRNAs is statistically indistinguishable from that of protein-coding genes (p=0.93, two-tailed Fisher’s exact test, Figure 2E), we cannot exclude this is in part a consequence of the lack of power to identify differences. Evidence that selection has purged deleterious mutations at elncRNA splicing-associated motifs during mammalian and recent human history, supports the functional relevance of elncRNA processing.

### Disruption of elncRNA splicing decreases target expression

We used genotyping data (The 1000 Genomes Project Consortium 2012) and gene expression data (Lappalainen et al., 2013) from LCLs derived from 373 healthy individuals to identify nucleotide variants that disrupt elncRNA splicing (Figure 3A). This unbiased approach allowed us to directly test whether changes in elncRNA splicing impact enhancer activity.

**Figure 3.**
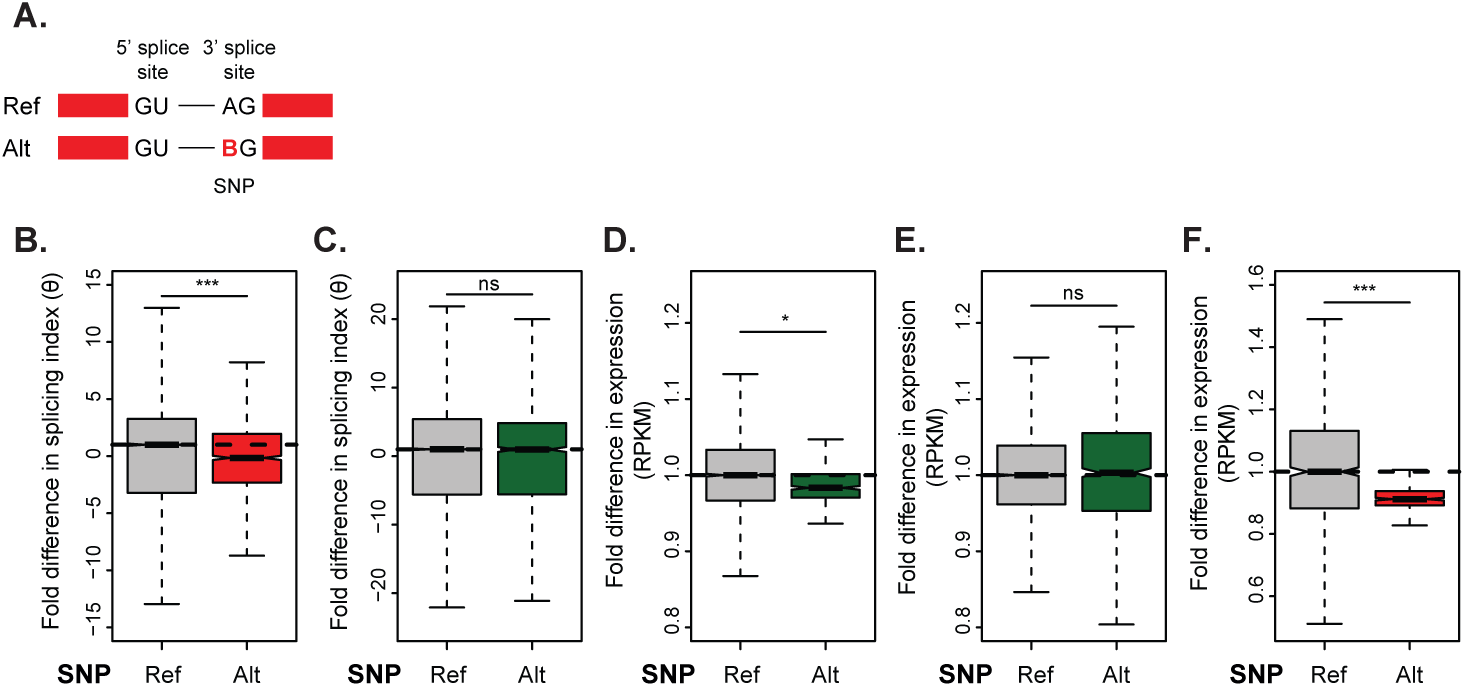
Disrupted elncRNA splicing impacts *cis*-gene regulation. (A) Example of a splice site variant that potentially disrupt elncRNA splicing. In contrast to individuals that carry the reference genome allele at a canonical 3’ splice site, those with alternative alleles have carry any nucleotide but A (denoted as B) at the first position of the acceptor site. Distribution of the median fold difference, between individuals that carry alternative or reference alleles at elncRNA splice sites, in splicing index of (B) multi-exonic elncRNAs and (C) target protein coding genes; (D) target and (E) non-target gene expression levels (RPKM); and (F) elncRNA expression. Differences between groups were tested using a two-tailed Mann-Whitney *U* test. * *p* < 0.05; *** *p* < 0.001; NS *p* > 0.05.

We identified 4 variants that disrupt the splice-donor or acceptor sites of 4 elncRNAs that have at least one predicted *cis-*target (Supplementary Table ST2, Methods). Since enhancer-promoter interactions rely on local chromosomal looping, enhancers and their associated transcripts often do not target the nearest gene in linear distance (Javierre et al. 2016). We unbiasedly predicted putative elncRNA targets as those that are jointly associated to the same expression quantitative trait loci (eQTL) variants (Methods), supporting their potential co-regulation (Tong et al. 2017).

We used an approach that explores spliced reads (intronic spanning reads) to quantify differential abundance of spliced isoforms in individuals with reference and alternative alleles at splice sites (Li et al. 2018). Individuals that carry nucleotide variants disrupting canonical splice donor/acceptor site have significantly decreased transcript splicing efficiency relative to individuals with the reference canonical splice sites allele (GT-AG) (Figure 3B, Supplementary Figure S3). As expected, the impact of SS variants on gene splicing efficiency depends on the total number of alternative transcripts and exons and ranges from 11% to 24% for elncRNA with 6 to 2 number of exons, respectively (Supplementary Figure S3). Consistent with a direct role of splicing in the modulation of enhancer function, this natural mutational study revealed that disruption of elncRNA splicing, which does not impact their protein-coding target splicing (p>0.3, two-tailed Mann-Whitney *U* test, Figure 3C, Supplementary Figure S3), is associated with significant decrease in target expression (p<0.05, two-tailed Mann-Whitney *U* test, Figure 3D, Supplementary Figure S3). The association between of splice site mutations and gene expression was restricted to predicted elncRNA targets genes, as the expression levels of other nearby was unaffected (p>0.1, two-tailed Mann-Whitney *U* test, Figure 3E, Supplementary Figure S3). In addition to changes in target protein-coding gene levels, the relative abundance of elncRNAs was also significantly decreased in individuals that carried variants that altered elncRNA splice donor or splice acceptor sites (Figure 3F, Supplementary Figure S3). This is consistent with the well-established synergy between RNA processing and transcription and the impact of co-transcriptional splicing on gene expression (Furger et al. 2002; Damgaard et al. 2008). We replicated this mutational study using 89 samples of Yoruba (YRI) population from the Geuvadis dataset (Supplementary Figure S4) and the analysis of these results consistently supports that elncRNA splicing variants impact cognate enhancer function.

### elncRNA splicing causally mediate target expression

The results of these mutational studies support the role of splicing in modulating enhancer activity and suggest that this is at least in part a consequence of increased transcription of multi-exonic elncRNAs. However, the interdependence between elncRNA splicing and expression together with the fact that splice site mutations may also affect enhancer factor binding make it difficult to directly use splice site mutations to disentangle the distinct roles of splicing and transcription on enhancer function. Furthermore, although unbiased, splice site mutations were only found for a subset of elncRNAs, restricting the extent of our analysis. To overcome these limitations, we first used multi-variate regression to investigate the relationship between splicing of elncRNAs and the expression of their putative targets genome-wide. First, we found that elncRNA splicing is correlated with target expression levels (Supplementary Figure S5A) and this correlation is significantly higher than compared to that with their proximal non-targets (p<2.2×10^−16^, two-tailed Mann-Whitney *U* test, Supplementary Figure S5B). Next, we found by adding elncRNA splicing to its expression significantly, albeit moderately, improved the predicative power of its target gene expression (p=0.02, two-tailed Mann-Whitney *U* test, Supplementary Figure S5C). This is in contrast to elncRNA proximal non-targets, whose abundance was not explained by elncRNA splicing (p=0.8, two-tailed Mann-Whitney *U* test, Supplementary Figure S5C), supporting that the impact of elncRNA splicing on gene expression is target-specific.

Next, we expanded the number of elncRNA loci considered in our mutational study by identifying SNPs associated with the amount of splicing at multi-exonic elncRNAs (sQTLs) (Figure 4A). To directly assess the impact of elncRNA splicing on enhancer activity independent of its transcription, we excluded sQTL variants that are also associated with elncRNA expression (eQTLs) from this analysis (Methods). The average distance between these variants and elncRNA transcriptional start sites and their associated-enhancer boundaries is kb and 72.2 kb, respectively (Supplementary Figure S6A-B), eliminating the potential confounding impact of sQTLs on enhancer factor binding. We identified sQTLs for 25% (88/352) of multi-exonic elncRNAs. Of these, more than half (49, 56%) had at least one sQTL that was also associated with their target expression (80 elncRNA-target pairs, 2,354 joint seQTLs).

**Figure 4.**
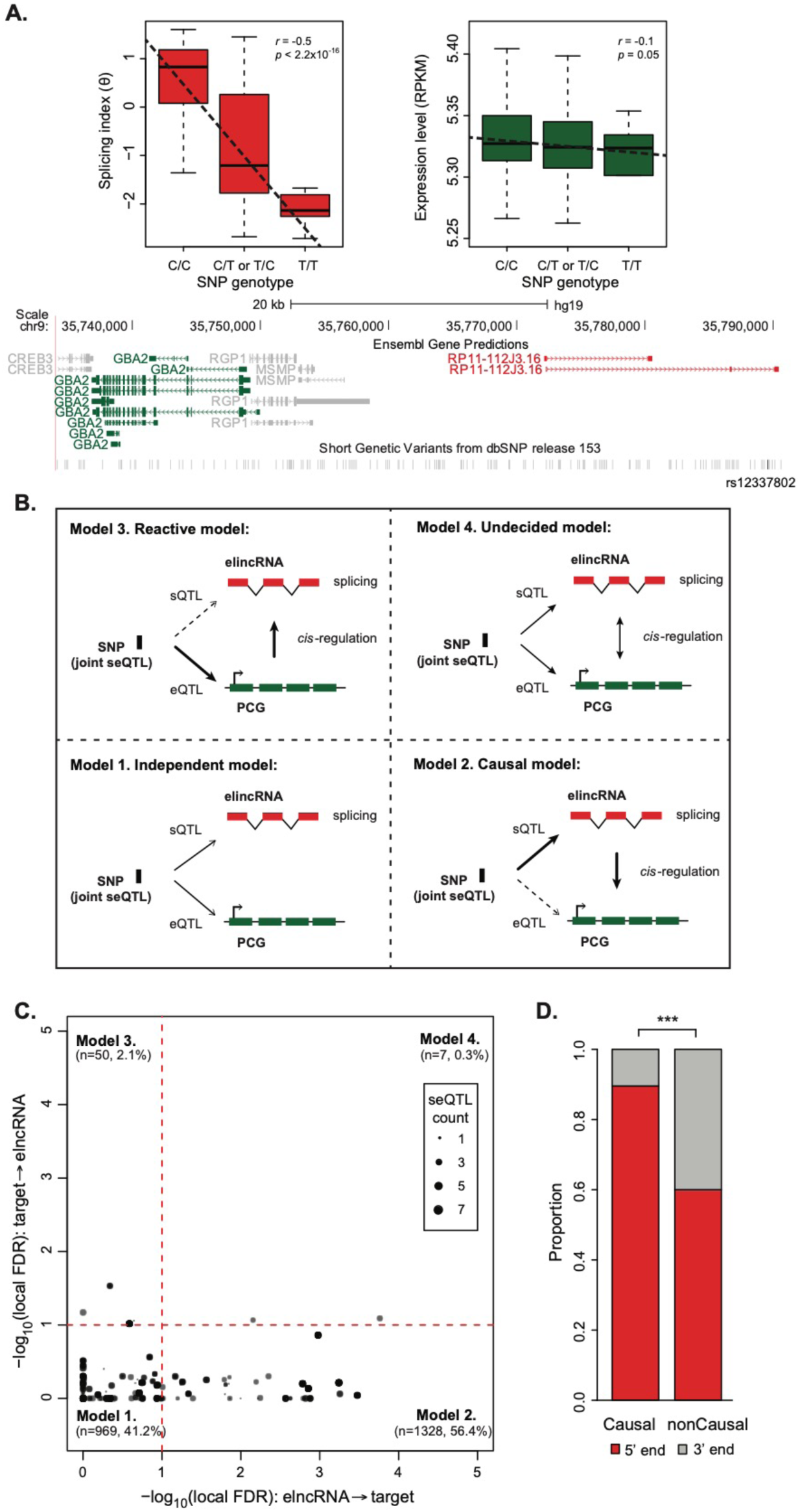
Impact of elncRNA splicing on *cis*-gene regulation in the human population. (A) RP11-112J3 (ENSG00000227388) is a multi-exonic elncRNA whose splicing is associated with genotype of the SNP variant (rs12337802), which is also associated to the expression level of an elncRNA target, GBA2 (ENSG00000070610). (Top panel) Distribution of the splicing index of elncRNA (RP11-112J3, red) and target expression (ENSG00000070610, green) in samples across the population that carry different alleles of the SNP variant (rs12337802). Spearman’s rho and p-values are shown. (Bottom panel) Genome browser illustrating the genomic positions of the elncRNA (RP11-112J3, red), its target (GBA2, green), and their associated SNP (rs12337802, black). (B) Four models of causal inference testing that predict the relationship between joint seQTL variant (black box) associations with the splicing (sQTL) of multi-exonic elncRNAs (red boxes) and the expression level (eQTL) of their target protein-coding genes (green boxes). Schematic representation of the models of joint seQTL associations: (1) the variants are independently associated with elncRNA splicing and target expression (independent model); (2) direct association between the variant and elncRNA splicing mediates the indirect association between that and target expression (causal model); (3) direct association between the variant with target expression mediates the indirect association between that and elncRNA splicing (reactive model); and (4) the causative interaction between elncRNA splicing and target expression is more complex (undecided model). Direct associations are depicted as solid lines and indirect associations as dash lines. (C) Scatterplot depicting causal inference testing local FDR associated with each the four models (as illustrated in B). Number and proportion of joint seQTLs are provided in brackets for each model. Dotted red lines denote significance threshold at local FDR < 0.1. (D) Proportion of joint elncRNA seQTLs causally or non-casually predicted to mediate target gene expression at the 5’ end (red) or 3’ end (grey) ends of the transcripts. Differences between groups were tested using a two-tailed Fisher’s exact test. *** *p* < 0.001.

Given that quantitative trait variants were identified using a relatively small cohort (373 LCL samples) and that cohort size is an important factor in QTL calling, we reasoned that the relatively small fraction of elncRNAs (∼14%, 49/352) with joint seQTL associations is a consequence of our study being under-powered. Using a dataset with nearly 85-times more samples (31,684 blood samples, eQTLGen consortium) (Võsa et al. 2018), we found nearly 80% of elncRNAs (70/88) splicing QTLs are jointly associated to target expression (171 elncRNA-target pairs). Compared to the smaller LCL data, target-seQTL associations were considerably higher (3.45×10^61^ times, p<2.2×10^−16^, two-tailed Mann-Whitney *U* test, Supplementary Figure S6C). In addition, using the larger dataset, target eQTLs that were not identified to be associated with elncRNA splicing by the smaller LCL set (non-seQTLs) were associated to a similar level as those identified to be jointly associated to elncRNA splicing (seQTLs) (p=0.3, two-tailed Mann-Whitney *U* test, Supplementary Figure S6D). This corroborates that our analysis is limited by the sample size of the LCL data and we are likely to be under-estimating the prevalence of elncRNAs whose splicing contribute to enhancer activity. However, 80% of the expression data used by the eQTLGen consortium was generated using microarray technology which cannot be used to predict splicing frequency. Given this limitation and the restricted access to its sensitive data, we could not carry out detailed analysis using the eQTLGen data and continued to use the LCL dataset.

Different models can account for the joint association between a variant and elncRNA splicing or its target expression (joint seQTL, Figure 4B) (Li et al. 2010). This can occur if the variant is independently associated with elncRNA splicing and target expression (independent model) or as the result of the complex interaction between splicing and expression (undecided model) Alternatively, variant can be associated with the target expression that in turn mediates splicing of elncRNA (reactive model). Finally, cases where the seQTL is associated with elncRNA splicing that in turn mediate target expression (causal model) would support a functional role of elncRNA splicing. For each seQTL, we established the most likely model of association using causal inference analysis (Wang and Michoel 2017). With a false discovery rate of 10%, we found that most associations between seQTLs (Figure 4C) and target expression is causally mediated by elncRNA splicing (n=1328, 56.4%). These causal seQTLs support the role of splicing for 61% of elncRNAs in the regulation of target expression (1 to 2 targets on average). Of the remaining variants, most are independently associated with elncRNA splicing and target expression (n=969, 41.2%), which is likely in part due to the relatively high false negative rates of this type of analysis (Wang and Michoel 2017). To account for the effect of linkage disequilibrium and noise in genotyping data, we repeated the analysis using only seQTLs that are the best (ie. most significant) variant associated with elncRNA splicing. Using this conservative set of seQTLs, we identified a similar proportion of elncRNAs (60%) whose splicing is predicted to causally regulate their target expression (Supplementary Figure S7), corroborating the causal impact of elncRNA splicing on their cognate enhancer function.

Importantly, while the fraction of seQTL associated with 5’ and 3’ end splice junctions is similar, 90% of those that support elncRNA splicing as a mediator of target expression are located at the 5’ end of the transcript, which is consistent with the known synergy between transcription and 5’ end splicing (Figure 4D).

### elncRNA splicing impacts local chromatin state

Evidence that variants associated with changes in elncRNA splicing also mediate target gene expression support the direct role of splicing on enhancer function. To assess if elncRNA splicing impacts cognate enhancer function by directly impinging on its activity, we investigated changes in chromatin signatures at elncRNA cognate enhancers. This was possible thanks to the recent release of genome-wide histone modification data, including H3K4me1 and H3K27ac, for a subset of 150 LCL samples (Delaneau et al. 2019). Consistent with a synergistic contribution of splicing to enhancer activity, individuals that carried nucleotide variants that altered elncRNA splice site also had significantly lowered cognate enhancer activity, which we estimated using levels of H3K4me1 and H3K27ac (Heintzman et al. 2009) at cognate enhancers (Figure 5A), supporting that splicing at elncRNA loci impacts local enhancer-associated epigenetic state.

**Figure 5.**
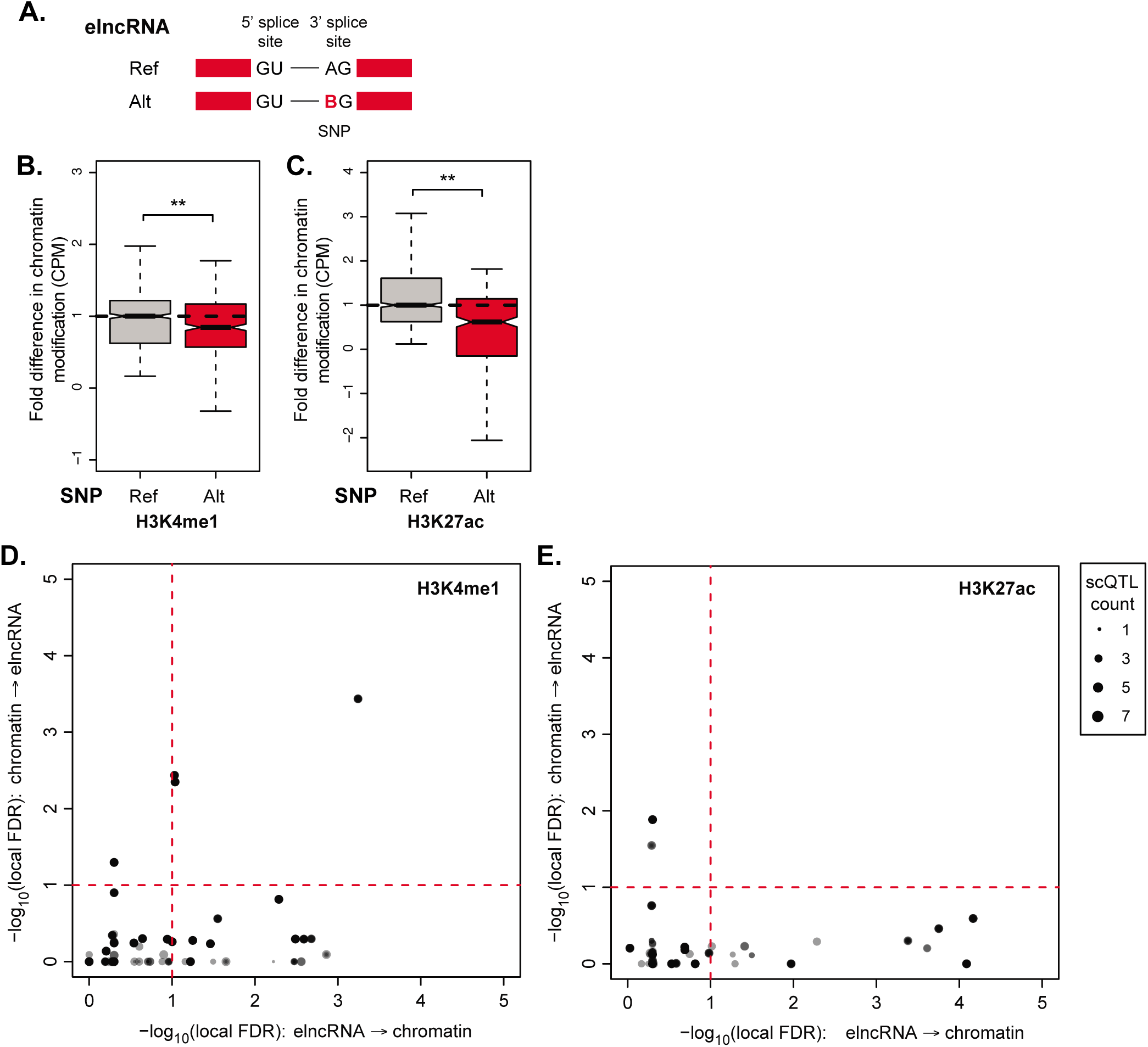
Impact of elncRNA splicing on cognate enhancer chromatin signatures in the human population. (A) Example of a splice site variant that potentially disrupt elncRNA splicing. In contrast to individuals that carry the reference genome allele at a canonical 3’ splice site, those with alternative alleles have carry any nucleotide but A (denoted as B) at the first position of the acceptor site. Distribution of the median fold difference in chromatin signature, between individuals that carry alternative or reference alleles at elncRNA splice sites, in (B) H3K4me1 and (C) H3K27ac. (D) Scatterplot depicting joint elncRNA scQTLs in which elncRNA splicing is causally or non-casually predicted to mediate enhancer chromatin signatures, (D) H3K4me1 and (E) H3K27ac, respectively, using causal inference testing as illustrated using local FDR associated with the four models (as shown in Figure 4B). Dotted black lines denote significance threshold at local FDR < 0.1. Differences between groups were tested using a two-tailed Mann-Whitney *U* test. ** *p* < 0.01.

To extend this analysis genome-wide, we next identified nucleotide variants associated with elncRNA splicing and chromatin modification (scQTL, 1,932 and 534 scQTLs involving 30 and 23 elncRNAs, for H3K4me1 and H3K27ac, respectively). We determined the most likely relationship between scQTLs, elncRNA splicing and chromatin signatures using causal inference analysis (Wang and Michoel 2017). We found that a large proportion of scQTL variants (855 (44.3%) and 397 (74.3%) scQTLs for H3K4me1 and H3K27ac, respectively) associated independently to elncRNA splicing and chromatin mark levels. This is expected given the relatively small number of samples available, which has likely reduced the power of this analysis as discussed above. Despite this limitation, we predicted that splicing of a sizable fraction of elncRNAs modulates chromatin state (40% and 48% for H3K4me1 and H3K27ac, respectively, Figure 5D,E). These fractions were significantly higher compared to the fraction of elncRNAs whose splicing were predicted to be impacted by chromatin modification changes (3% and 4% for H3K4me1 and H3K27ac, respectively; p<0.05, two-tailed Fisher’s exact test, Figure 5D,E). We conclude that splicing a sizable fraction of enhancer-associate transcripts directly impacts the activity of their cognate enhancer activity.

## DISCUSSION

We and others have previously shown that splicing of multi-exonic elncRNAs is strongly associated with high enhancer activity in several human and mouse cell lines (Gil and Ulitsky 2018; Tan et al. 2020). The direct contribution of splicing to enhancer activity has been shown for a few candidate elncRNA transcripts (Yin et al. 2015; Engreitz et al. 2016). However, it remains uninvestigated whether elevated enhancer activity resulting from the splicing of these elncRNAs is a general phenomenon that would explain their previously described genome-wide associations (Gil and Ulitsky 2018; Tan et al. 2020).

To gain initial insights into this, we first investigated the evolution of splicing-associated motifs, including splice sites, within elncRNAs. We found sequences that support elncRNA splicing evolved under purifying selection, an evolutionary signature of functionality. Interestingly, splicing related motifs were previously found to be amongst the most highly conserved sequences within lncRNAs (Haerty and Ponting 2015). With at least half of all lncRNA transcripts initiating at enhancers (Marques et al. 2013; Hon et al. 2017) and a sizable proportion of these undergoing splicing (Gil and Ulitsky 2018; Tan et al. 2020), it is likely the relative high sequence constraint observed at splicing-associated motifs within lncRNAs (Haerty and Ponting 2015) can be attributed to enhancer transcribed lncRNAs. Consistent with this, we found splicing-related motifs within the less efficiently processed promoter-associated lncRNAs (plncRNAs) (Tan et al. 2020) to be less constrained.

We investigated the direct contribution of elncRNA splicing to cognate enhancer activity using a statistical population genomics approach in human lymphoblastoid cell lines (LCLs). We found splicing of at least 60% of elncRNAs causally mediates changes in their target gene expression levels. Interestingly, a sizable proportion (40%) of elncRNA splicing was also predicted to mediate changes in chromatin signatures, including H3K4me1 and H3K27ac, at enhancers. The present analysis is limited by the size of the cohort analysed. Relatively small sample sizes reduce the ability to confidently identify quantitative trait loci (Beavis 1998). This is illustrated by 1.4-fold increase in the number of seQTL detected when using data from an much larger cohort (eQTLGen (Võsa et al. 2018)) than the one from Geuvadis (Lappalainen et al. 2013). We anticipate that the use of larger cohorts should result in an increase in the percentage of elncRNAs for which splicing can be confidently shown to mediate target expression in *cis*.

Our analysis of elncRNA splice-motif evolution and the impact of elncRNA processing on enhancer function and activity support that elncRNA splicing is unlikely a spurious by-product of enhancer activity. This is consistent with evidence supporting conserved elncRNA splicing and their lack of exonic constraint (Haerty and Ponting 2013; Tan et al. 2020), suggesting the contribution of elncRNA splicing to the modulation of enhancer activity go beyond the processing of their mature transcripts.

Whereas the precise molecular mechanisms underlying this association remain currently unknown, a number of mechanisms can support the role of elncRNA splicing on enhancer activity. Splicing has long been known to directly impact the rate of transcription (Le Hir et al. 2003). For example, DNA elements embedded within introns have been shown to contribute to transcriptional regulation (Sleckman et al. 1996) and components of the spliceosome can directly enhance RNAPII initiation (Kwek et al. 2002) and transcript elongation (Fong and Zhou 2001). Furthermore, it was recently shown that novel exon splicing events can result in transcription initiation at novel exon proximal promoters, possibly through the recruitment of transcription machinery by splicing factors (Fiszbein et al. 2019).

The synergy between splicing and transcription may also be influenced by changes in local chromatin environment. For example, splicing pattern changes have been shown to correlate with histone modification and factors that alter chromatin structure, including chromatin signatures at active enhancer regions (Luco et al. 2010). On the other hand, tethering of enhancer-associated transcripts at their site of transcription has been shown to alter local chromatin environment (Lai et al. 2013; Li et al. 2013; Hsieh et al. 2014; Bose et al. 2017). Chromatin-bound lncRNAs have been recently shown to be enriched in U1 small nuclear ribonucleoprotein (snRNP) binding, a protein essential for the recognition of nascent RNA 5’ splice site and assembly of the spliceosome (Yin et al. 2020). The depletion of U1 snRNP reduced the tethering of lncRNAs to chromatin, suggesting a mechanism by which elncRNA splicing can strengthen enhancer activity (Yin et al. 2020).

The link between splicing and transcription, together with evidence of RNA-dependent recruitment and activity of some enhancer factors, provide initial clues to what might be the molecular mechanism(s) underlying the role of elncRNA splicing in the modulation of enhancer activity, as supported by the present work.

## METHODS

### Identification of enhancer-associated transcripts

We defined enhancer-associated transcripts by considering all ENCODE intergenic enhancers in human GM12878 lymphoblastoid cell line (LCL. 98,529 enhancers)(Bogu et al. 2015) overlapping DNase I hypersensitive sites (Encode Project Consortium 2012) and a CAGE cluster (Andersson et al. 2014) in LCLs (n=4997). We considered all LCL-expressed lncRNAs (Tan et al. 2017) and Ensembl annotated protein coding genes (version 75). LncRNAs whose 5’ end is within 500bp of a LCL enhancer were considered as being transcribed from the enhancer (n=564 elncRNAs, Supplementary Table ST1), and all remaining enhancers were presumed to transcribe eRNAs (n=4433). Transcripts within 500bp of ENCODE LCL promoters (Encode Project Consortium 2012) were classified as protein-coding genes (n=12,070) or promoter-associated lncRNAs (plncRNAs, n=600) depending on their biotype annotation. We found 62.4% of elncRNAs (n=352), 60.5% of plncRNAs (n=363), and 94.4% of protein-coding genes (n=11,392) to be multi-exonic.

### Read mapping and quantification

For all downloaded data sets, adaptor sequences were first removed from sequencing reads with Trimmomatic (version 0.33) (Bolger et al. 2014) and then aligned to the human reference genome (hg19) using HISAT2 (version 2.0.2) (Kim et al. 2015). Since available Geuvadis RNA sequencing and genotype data, LCL CAGE data and enhancer predictions, were all originally mapped to GRCh37 (hg19), our analyses were performed using the same human genome assembly.

### Metagene analysis of elncRNAs

Direction of transcription was assessed with metagene profiles of CAGE reads centered at LCL enhancers and gene transcriptional start sites (TSS) plotted using NGSplot (version 2.4) (Shen et al. 2014) as in (Tan et al. 2020). Enrichment of histone modifications, transcription factor binding, and gene expression levels were assessed using publicly available GM12878 DNase-seq, ChIP-seq and RNA-seq data sets (Supplementary Table ST3). Metagene profiles of sequencing reads centered at gene TSSs were visualized using HOMER (version 4.7) (Heinz et al. 2010).

### Analysis of chromosomal architecture

Enrichment of enhancer-associated transcripts at LCL loop anchors (Rao et al. 2014), relative to expectation, was assessed using the Genome Association Tester (GAT, version 1.3.6) (Heger et al. 2013). Specifically, loop positional enrichment was compared to a null distribution obtained by randomly sampling 10,000 times (with replacement) segments of the same length and matching GC content as the tested loci within mappable intergenic regions (as predicted by ENCODE (Hoffman et al. 2013)). To control for potential confounding variables that correlate with GC content, such as gene density, the genome was divided into segments of 10 kb and assigned to eight isochore bins in the enrichment analysis.

The frequency of chromosomal interactions within topologically associating domains (TADs) was calculated using LCL Hi-C contact matrices (Rao et al. 2014), as previously described (Tan et al. 2017).

### Identification of splicing-associated motifs

Density of human exonic splicing enhancers (ESEs), using a set of 238 computationally predicted motifs (Fairbrother et al. 2002) and a more stringent set of 54 motifs identified across multiple studies (Caceres and Hurst 2013), were predicted within LCL transcripts as previously described (Haerty and Ponting 2015). Specifically, exonic nucleotides (50 nt) flanking splice sites (SS) of internal transcript exons (> 100 nt) were considered in the analysis, after masking the 5 nt immediately adjacent to SS to avoid splice site-associated nucleotide composition bias (Fairbrother et al. 2002; Yeo and Burge 2004). Canonical U1 sites (GGUAAG, GGUGAG, GUGAGU) adjacent to 5’ splice sites (3 exonic nt and 6 intronic nt flanking the 5’ SS) were predicted as previously described (Almada et al. 2013). FIMO (MEME version 4.12) (Grant et al. 2011) was used to search for perfect hexamer matches within these sequences.

### GC content

For all LCL expressed genes with at least 2 exons, we computed the GC content for the first exon, remaining exons, and introns for each gene separately, as well as their flanking intergenic sequences of the same length (after excluding the 500 nucleotides immediately adjacent to annotations).

### Splicing efficiency

Transcript splicing efficiency was estimated by computing the proportion of fully excised introns using bam2ssj (IPSA version 3.3) (Pervouchine et al. 2013) of transcripts for each gene. Using long non polyA-selected RNA-seq data in GM12878 (Encode Project Consortium 2012), we calculated the splicing index, coSI (θ), which represents the ratio of reads that span exon-exon splice junctions (excised intron) over those that overlap exon-intron junctions (incomplete excision) (Tilgner et al. 2012).

### elncRNA conservation across evolution and in humans

To assess selective constraints of transcripts, we estimated their pairwise nucleotide substitution rates. First, pairwise alignment between human and mouse of the different splicing-associated features (including canonical splice sites, exonic splicing enhancers and canonical U1 sites) were separately concatenated for multi-exonic elncRNAs, plncRNAs and protein-coding genes. We used BASEML from the PAML package [version 4.9, REV substitution model (Yang 1997)] to estimate pairwise nucleotide substitution rates of each splicing motif. To determine whether the splicing motifs have evolved under significant purifying selection, we compared the observed nucleotide substitution rate to that of randomly selected pseudo-splicing motif sequences of the same length and GC content from neighbouring (within 1Mb) neutrally evolving sequences ancestral repeats (ARs, (Lunter et al. 2006)). We repeated this process 1000 times to obtain a distribution of the expected substitution rates.

To assess the conservation of splicing motifs during modern human evolution, we determined their derived allele frequency (DAF), as previously described (Haerty and Ponting 2013). We identified common single nucleotide polymorphisms (SNPs) in the human population (dbSNP build 150) mapped within splicing motifs of elncRNAs, plncRNAs and protein-coding genes. DAF was calculated using information on ancestral allele and the frequency of these SNPs in the European population obtained from the 1000 Genomes Project (The 1000 Genomes Project Consortium 2012). To determine the expected background distribution, we compared DAF spectrum of the observed splicing motifs to that of randomly selected sequences of the same length and GC content from local ARs.

### Mapping of molecular quantitative trait loci (QTLs)

Expression values (RPKM) of multi-exonic elncRNAs and protein-coding genes in EBV-transformed LCLs derived from 373 individuals of European descent (CEU, GBR, FIN and TSI) were quantified (as described in (Tan et al. 2017)). The corresponding processed genotypes were downloaded from EBI ArrayExpress (accession E-GEUV-1) (Lappalainen et al. 2013). Quantification of alternative splicing events was estimated using LeafCutter (version 1.0) (Li et al. 2018). Single nucleotide polymorphisms (SNPs) located within the same TAD as the genes of interest were tested for association with splicing (sQTLs) and with expression levels (eQTLs) of elncRNAs and protein-coding genes. Only SNPs with minor allele frequency (MAF) greater than 5% were considered in the QTL analyses. sQTLs and eQTLs were estimated using FastQTL (version 2.184) (Ongen et al. 2016). To assess the significance of the correlation globally, we permuted the splicing or expression levels of each gene 1000 times and noted the maximum permuted absolute regression coefficient (r_max_). We considered only sQTLs or eQTLs with an observed absolute regression coefficient (r_obs_) greater than 95% of all permuted r_max_ values to be significant (Lappalainen et al. 2013). We further performed Benjamini-Hochberg multiple testing correction to estimate FDR (<5%) for all SNPs within the same TAD. Putative protein-coding gene targets of multi-exonic elncRNAs were predicted as those that reside within the same TADs and whose expression levels were associated to the same SNP variant as the expression of the elncRNAs (ie. associated to the same eQTL). We found 88 elncRNAs to have putative protein-coding gene targets and are associated with at least one sQTL (n=26,741).

Levels of histone modification marks (H3K4me1 and H3K27ac, CPM) that overlap enhancers associated with elncRNAs were downloaded for the 150 European individuals with available chromatin data (Delaneau et al. 2019). To assess the relationship between elncRNA splicing and local histone modifications that mark enhancer elements, we calculated chromatin QTLs (cQTLs) associated with cognate enhancers of elncRNAs in the same way as sQTLs and eQTLs.

### Impact of genetic variation at elncRNA splice sites on cis-gene expression

We considered all SNPs located at elncRNA splice sites and estimated the fold difference in elncRNA splicing and steady state abundance, as well as in their putative target expression and chromatin marks at their cognate enhancer elements, between individuals that carry the reference or alternative alleles of these variants (Supplementary Table ST2).

### Causality inference between elncRNA splicing and nearby protein-coding gene expression

To infer the causal relationship between elncRNA splicing and putative target gene expression, we focused on QTLs that are associated with both splicing of elncRNAs (sQTL) and their putative target gene abundance (eQTL), and we refer to these variants as joint seQTLs. elncRNA sQTL variants that were also associated with elncRNA expression level or splicing of their putative *cis*-target genes were excluded from the analysis (n=21,650 out of 26,741). In total, we found 49 elncRNAs with seQTLs shared with 80 target genes.

For all triplets of seQTL – elncRNA splicing – target gene expression (n=2,349), we performed causal inference testing using a Bayesian approach as implemented by Findr (Wang and Michoel 2017) by testing the models: (1) the independent model where seQTL variants are independently associated with elncRNA splicing and target gene expression; (2) the causal model with elncRNA splicing as the molecular mediator of gene expression; (3) the reactive model where gene expression mediates elncRNA splicing; and (4) the undecided model where causative interaction between elncRNA splicing and target gene expression is more complex (Li et al. 2010).

The same causal inference testing was performing between elncRNA splicing (sQTL) and chromatin marks (cQTL) at their cognate enhancers by identifying scQTLs. We found 30 and 23 elncRNAs whose splicing-QTL is also associated with H3K4me1 (n=1391 triplets) and H3K27ac (n=532 triplets) marks, respectively, at their cognate enhancers.

### Linear regression

We used multi-variable regression to assess whether elncRNA splicing contributes to their cognate enhancer activity by comparing the difference in the proportion of variance in target expression is explained by adding elncRNA splicing to the model (Target-expression ∼ elncRNA-expression compared to Target-expression ∼ elncRNA-expression + elncRNA-splicing).

### Statistical tests

All statistical analyses were performed using the R software environment for statistical computing and graphics (R Development Core Team 2008).

### Data Access

Analyses were performed using publicly available command-line tools. Versions and deviations from parameters used are as detailed in the Methods. All scripts used to parse the results are available upon request.

## Acknowledgements

We thank members of the Marques group for valuable comments and discussion. We thank Zoltán Kutalik and Diogo Ribeiro for discussion on population genomics analysis; Olivier Delanau for early access to chromatin mark quantification. This work is funded by the Swiss National Science Foundation grant (PP00P3_179065 to ACM).

## Authors’ contributions

JYT and ACM designed the study. JYT performed the experiments and analysed the results. JYT and ACM discussed the results. ACM supervised the study. JYT and ACM wrote the manuscript. All authors approved the manuscript.

---

**Supplementary Figure S1.**
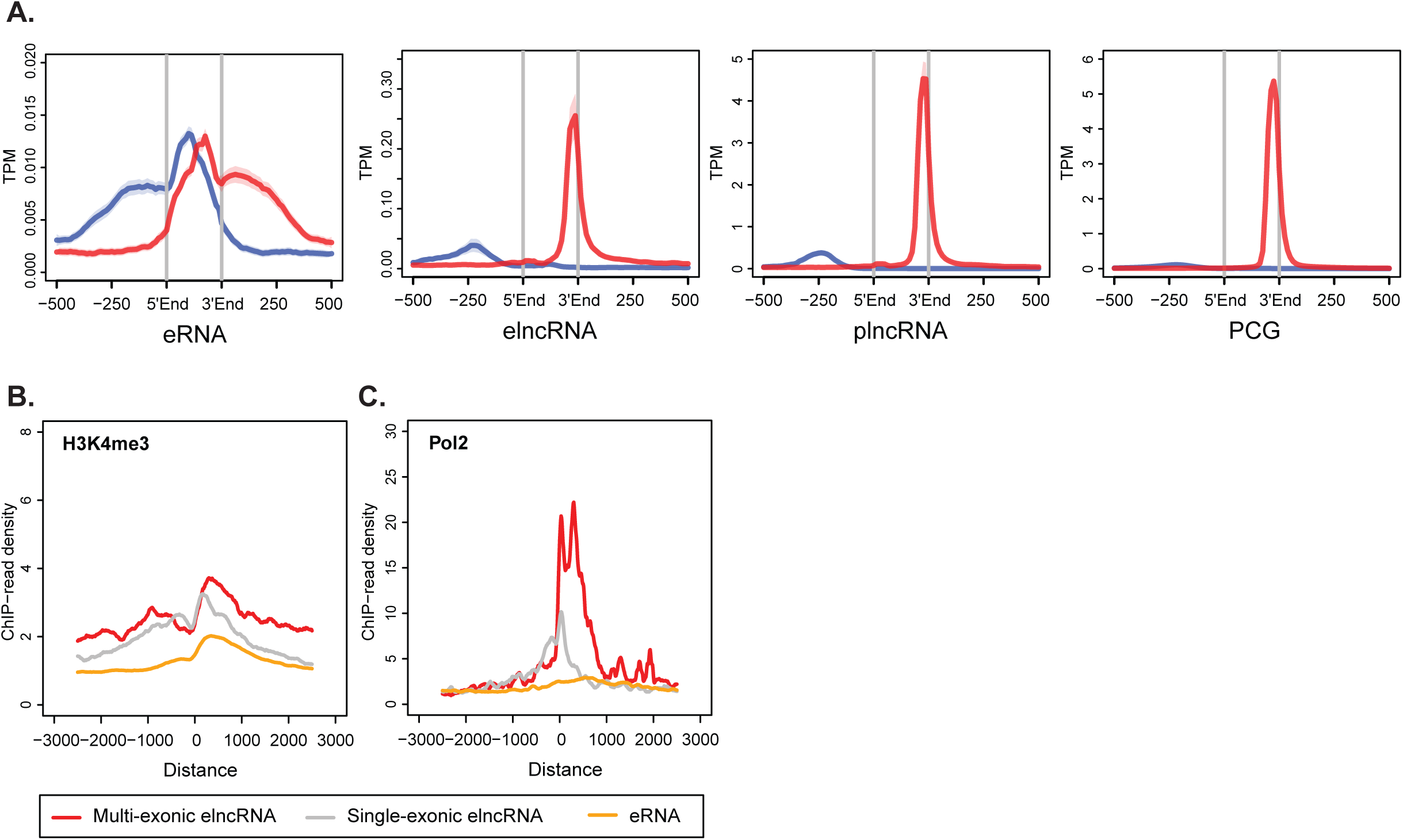

**Supplementary Figure S2.**
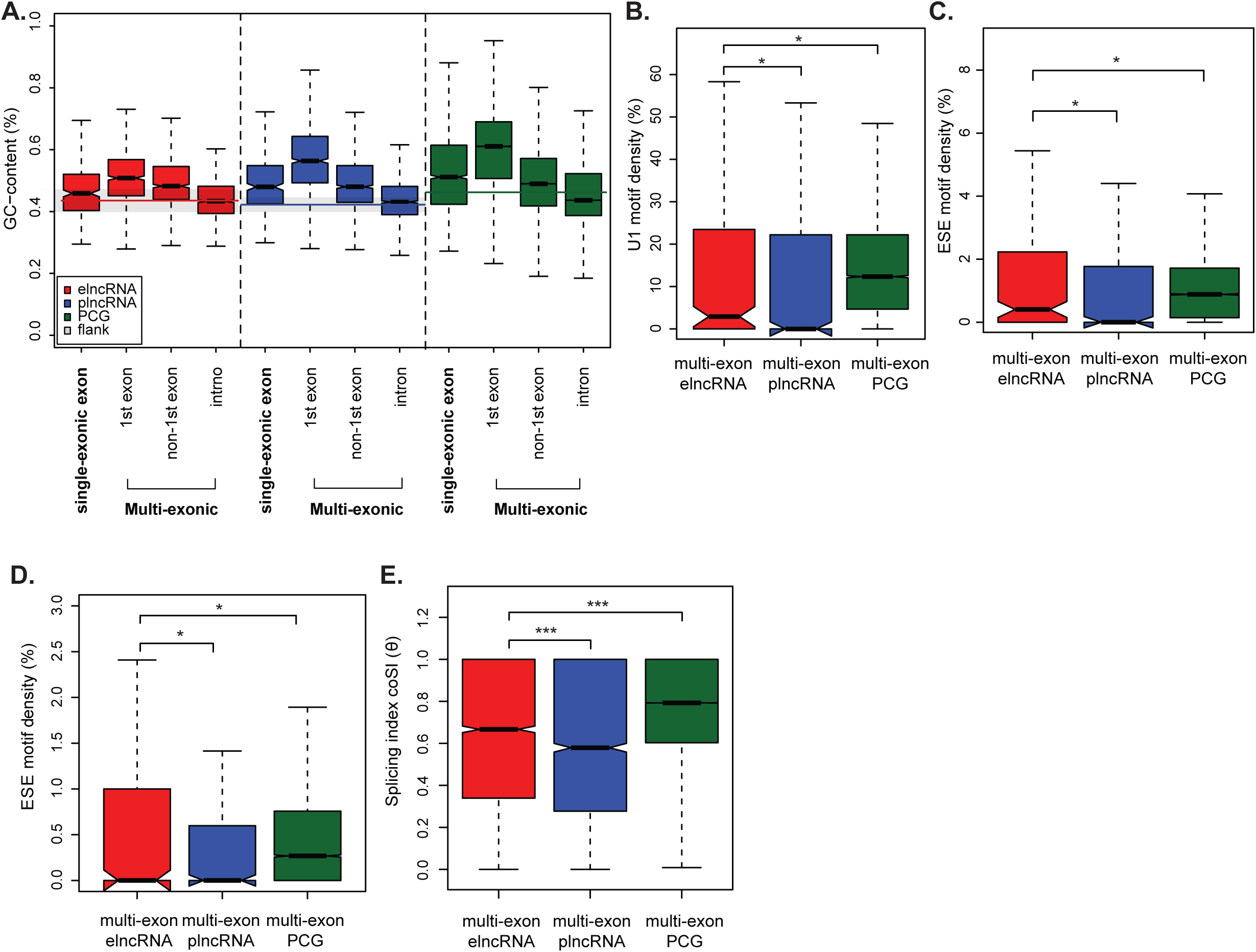

**Supplementary Figure S3.**
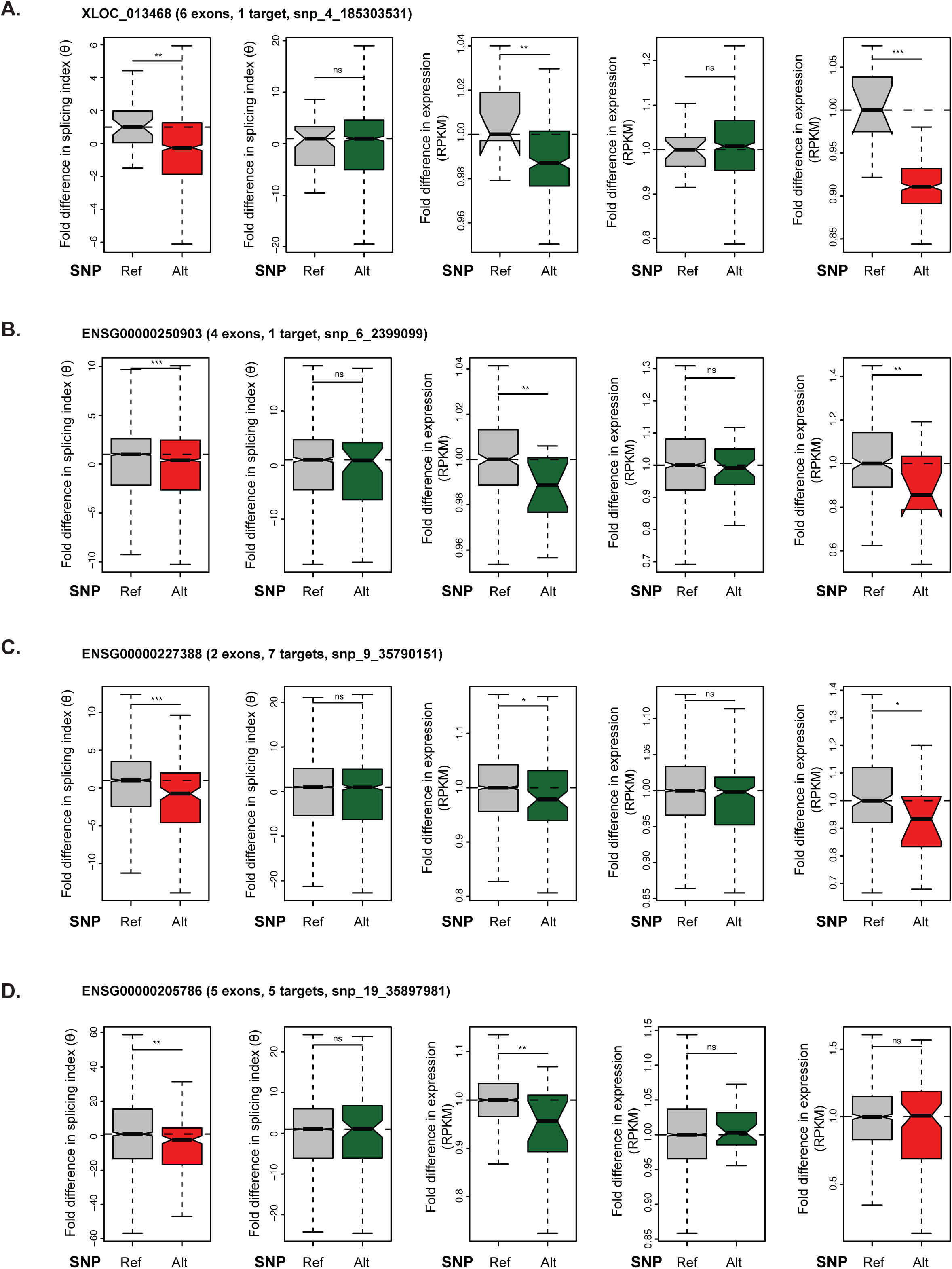

**Supplementary Figure S4.**
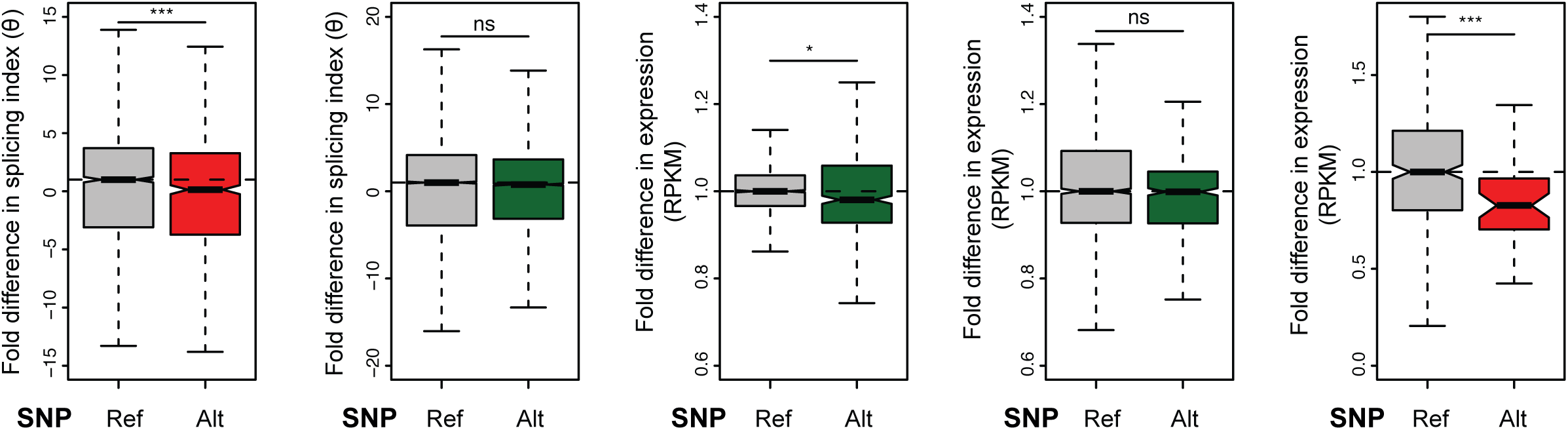

**Supplementary Figure S5.**
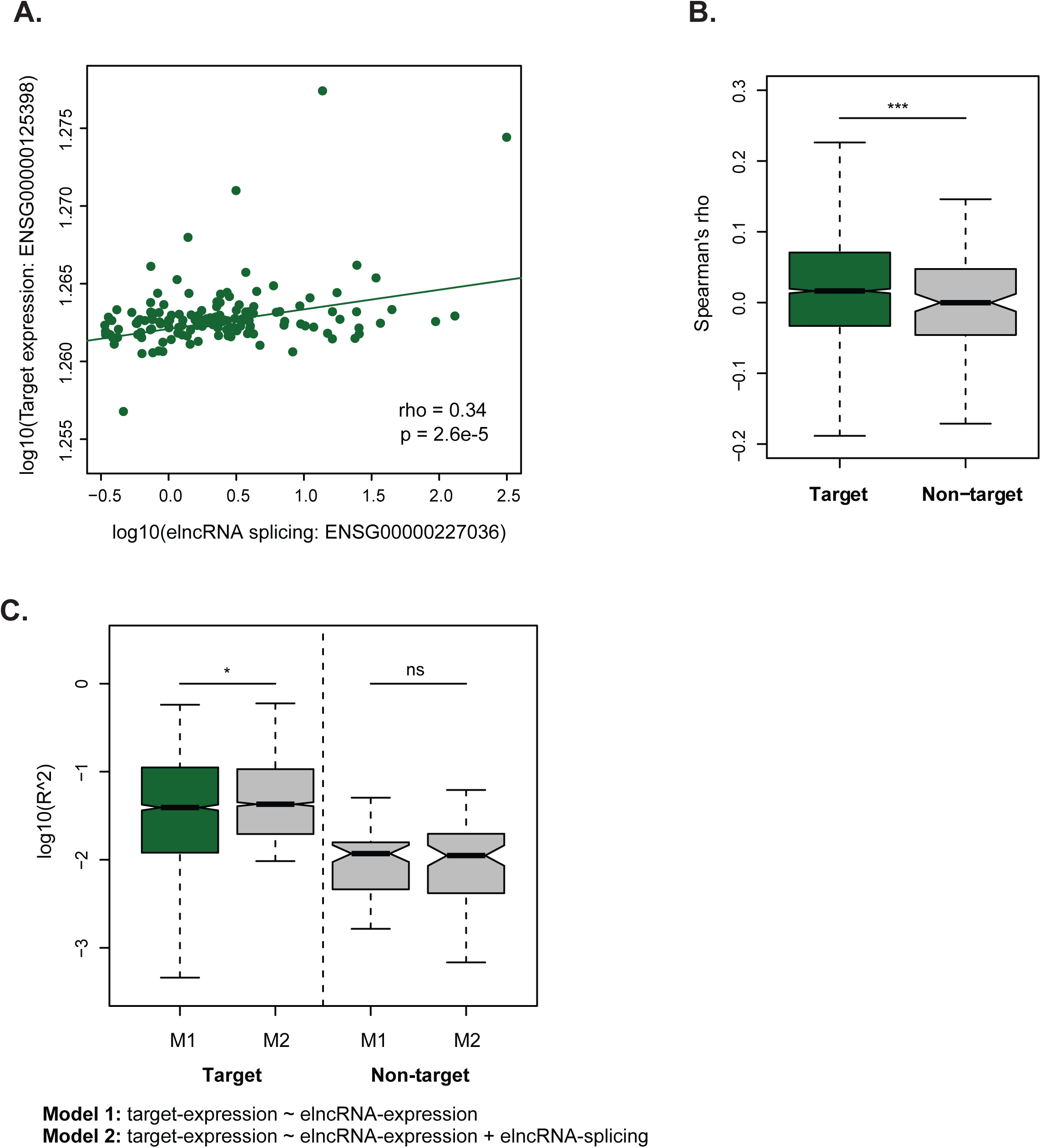

**Supplementary Figure S6.**
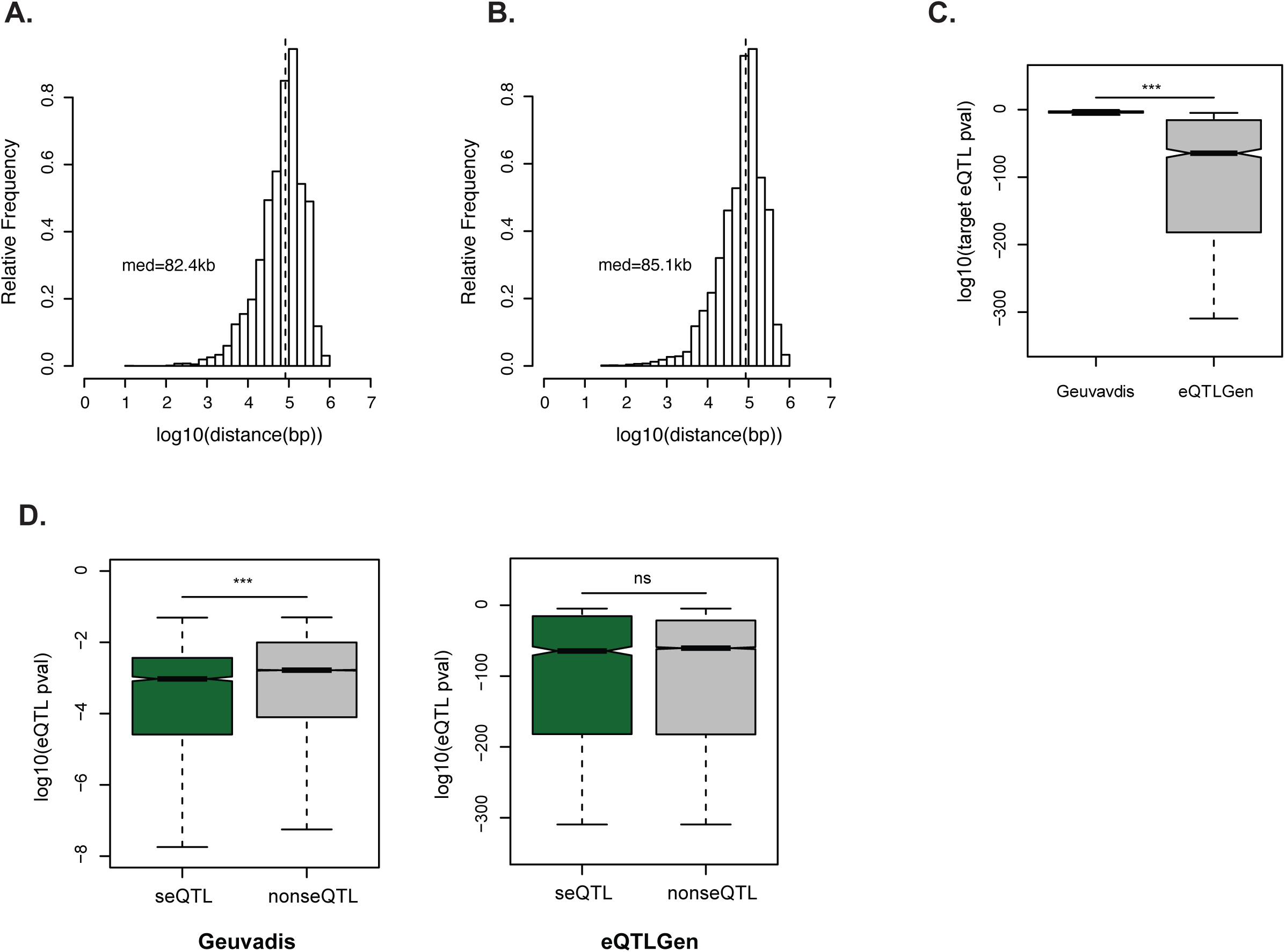

**Supplementary Figure S7.**
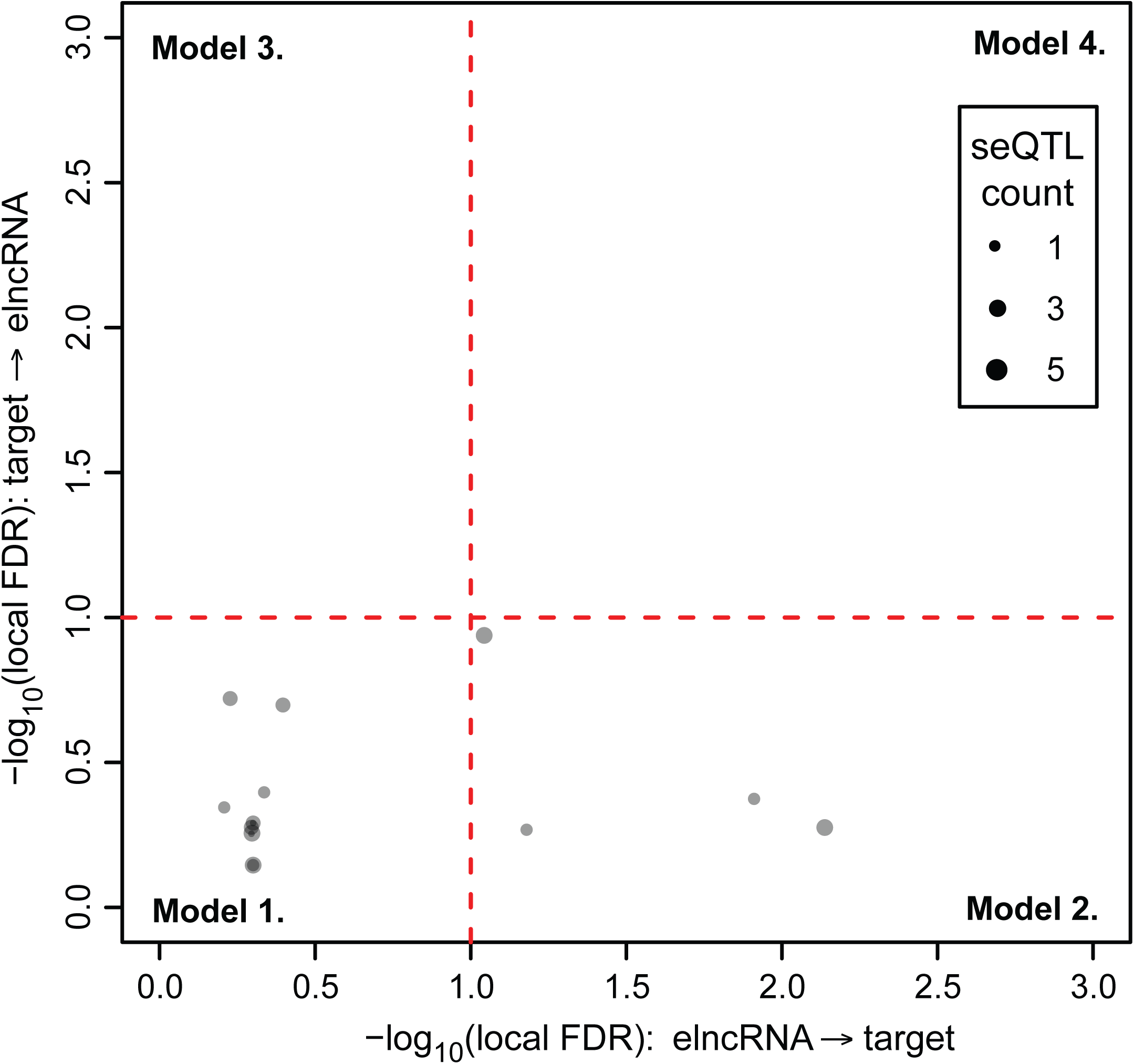

