## Supplementary_notes for "The activity of human enhancers is modulated by the splicing of their associated lncRNAs"

### SUPPLEMENTARY MATERIALS

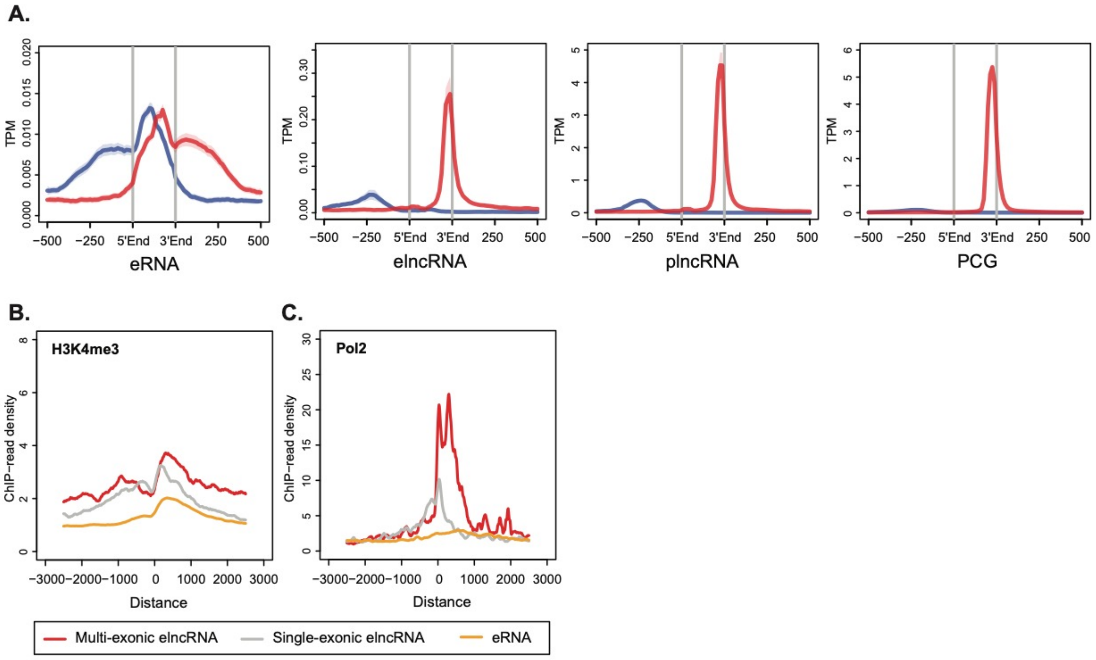


**Supplementary Figure 1. Multi-exonic elncRNAs are transcribed from highly active enhancers.** (A) Metagene plots of CAGE reads centered at transcription initiation regions (TIRs) of eRNAs, and promoters (estimated as -500bp to annotated gene TSS) of elncRNAs, plncRNAs and protein-coding genes (PCGs). Sense (red) and antisense (blue) reads denote those that map to the same or opposite strand, respectively, as the direction of their cognate TIRs. Metagene plots of (B) H3K4me3, (B) RNA Polymerse II ChIP-seq reads in LCLs at promoters of multi-exonic (red) and single-exonic (grey) elncRNAs, and eRNAs (yellow).

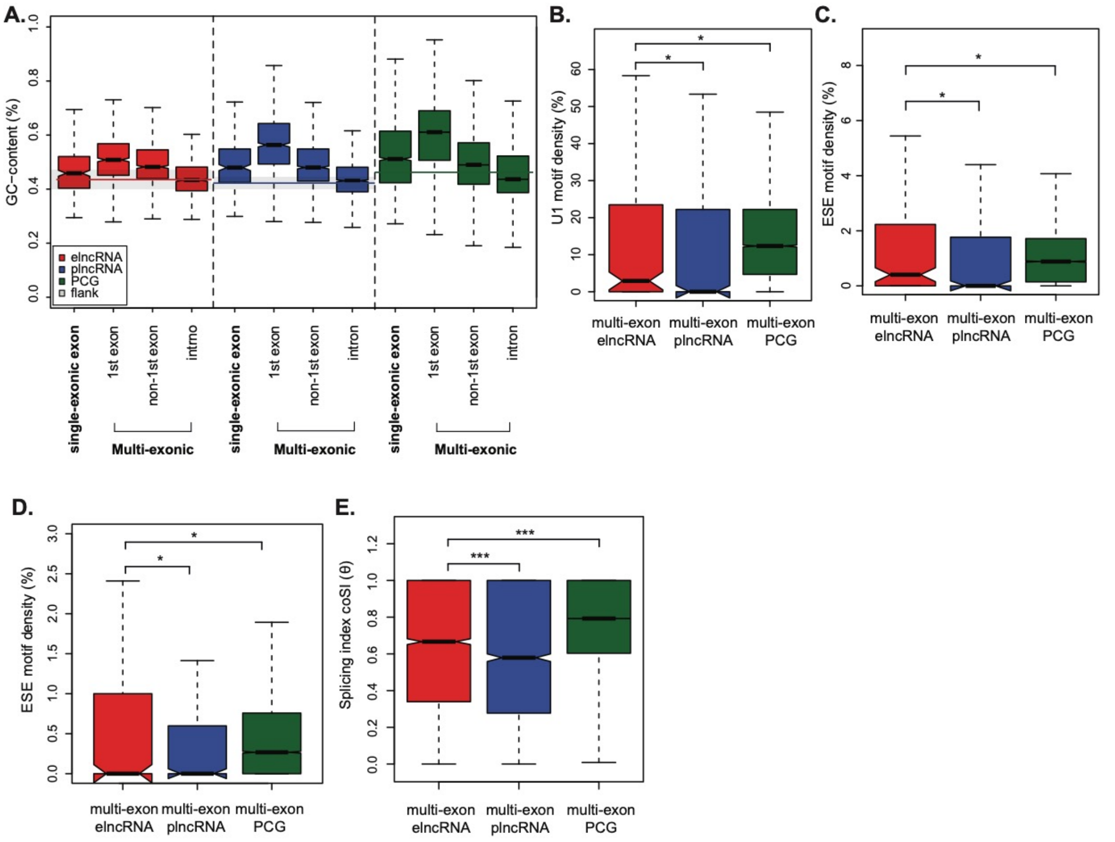


**Supplementary Figure 2. Splicing of elncRNAs has evolved under purifying selection.** (A) Distribution of GC-content across exons and introns of elncRNAs (red), plncRNAs (blue), and protein-coding genes (green). Distribution of the density of predicted (B) U1 spliceosome RNAs (snRNPs) and exonic splicing enhancers (ESEs), using (C) a comprehensive (n=238) and (D) a smaller but more stringent set (n=54) of predicted ESE motifs, within multi-exonic elncRNAs (red), plncRNAs (blue) and protein-coding genes (green). (E) Distribution of the splicing index, coSI ($\theta$) for multi-exonic elncRNAs (red), plncRNAs (blue) and protein-coding genes (green). Differences between groups were tested using a two-tailed Mann-Whitney *U* test. * $p < 0.05$; ** $p < 0.01$; *** $p < 0.001$; NS $p > 0.05$.

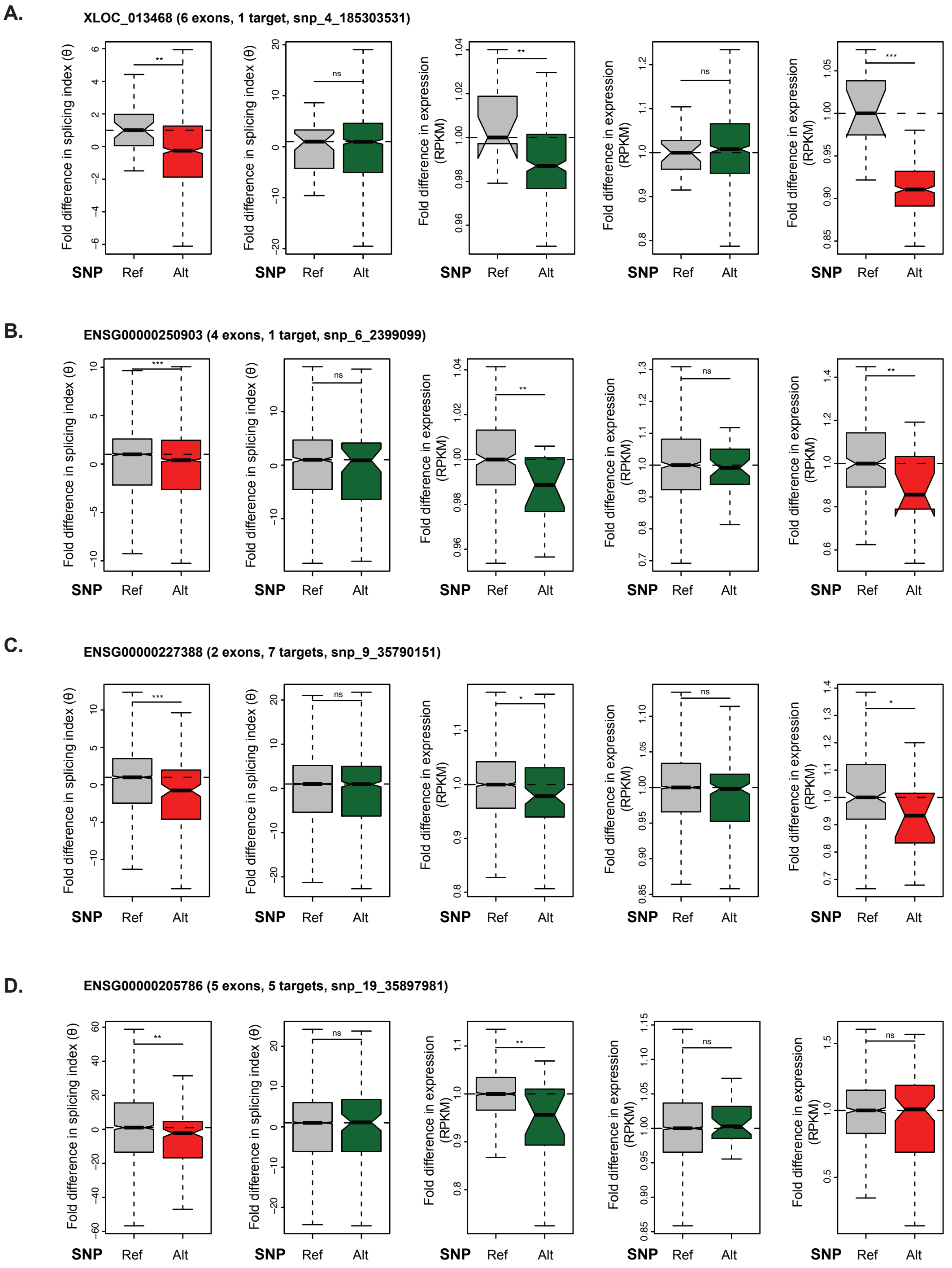


**Supplementary Figure 3. Disrupted elncRNA splicing impacts *cis*-gene regulation.** Distribution of the median fold difference, between individuals that carry alternative or reference alleles at splice sites of (A) XLOC_013468, (B) ENSG00000250903, (C) ENSG00000227388, and (D) ENSG00000205786, in splicing index of the multi-exonic elncRNA and its target protein coding genes; its target and non-target gene expression levels (RPKM); and elncRNA expression. Differences between groups were tested using a two-tailed Mann-Whitney *U* test. * $p < 0.05$; *** $p < 0.001$; NS $p > 0.05$. * $p < 0.05$; ** $p < 0.01$; *** $p < 0.001$; NS $p > 0.05$.

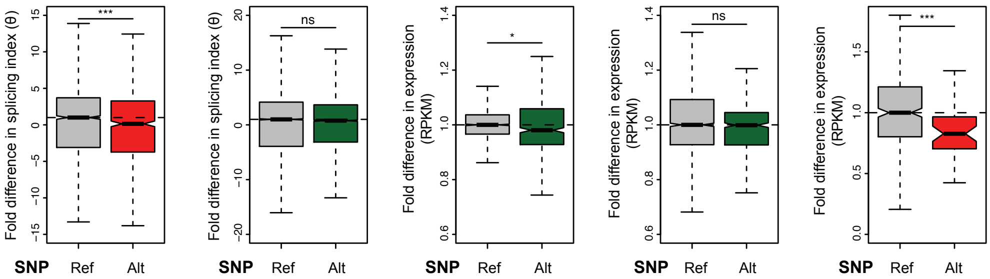


**Supplementary Figure 4. Disrupted elncRNA splicing impacts *cis*-gene regulation across YRI population.** Distribution of the median fold difference, between individuals of the Yorubin (YRI) population that carry alternative or reference alleles at elncRNA splice sites, in splicing index of multi-exonic elncRNAs and target protein coding genes; target and non-target gene expression levels (RPKM); and elncRNA expression. Differences between groups were tested using a two-tailed Mann-Whitney *U* test. * $p < 0.05$; ** $p < 0.01$; *** $p < 0.001$; NS $p > 0.05$.

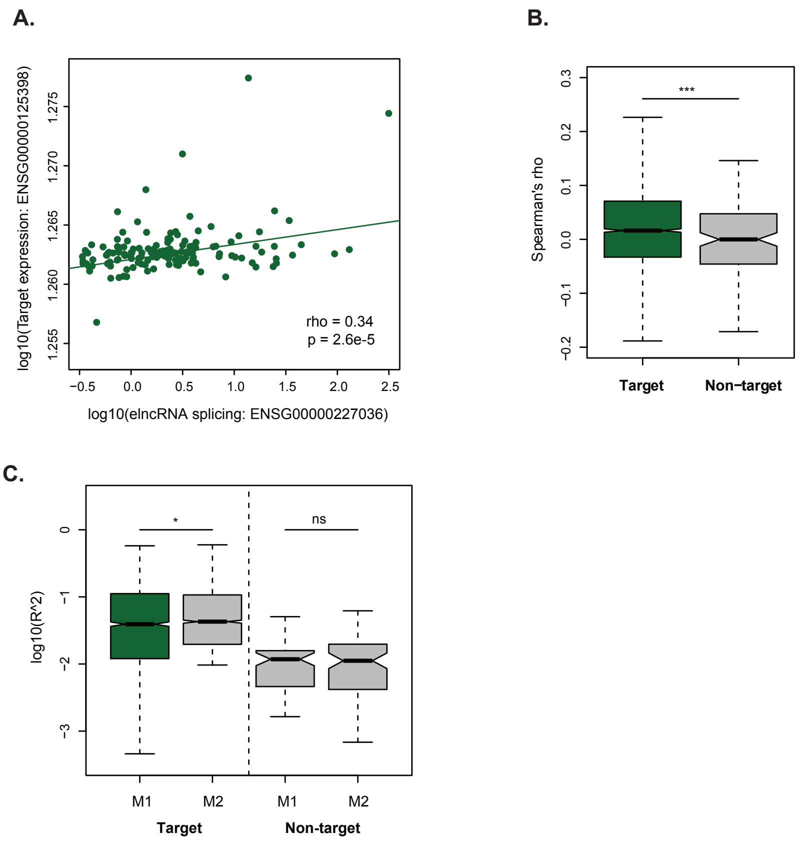


**Model 1:** target-expression ~ elncRNA-expression
**Model 2:** target-expression ~ elncRNA-expression + elncRNA-splicing

**Supplementary Figure 5. elincRNA splicing contributes to target expression levels.** (A) elncRNA splicing is significantly correlated (Spearman's test) with their *cis*-target expression levels, as illustrated using ENSG00000227036 (elncRNA) and ENSG00000125398 (target protein-coding gene). (B) Distribution of Spearman's correlation between elncRNA splicing and either their targets or non-targets. (C) Distribution of adjusted R square comparing between two regression models that use elncRNA expression and its splicing to predict expression of their targets or non-targets: Models 1: PCG-expression ~ elncRNA-expression; Model 2: PCG-expression ~ elncRNA-expression + elncRNA-splicing. Differences between groups were tested using a two-tailed Mann-Whitney *U* test. * $p < 0.05$; ** $p < 0.01$; *** $p < 0.001$; NS $p > 0.05$.

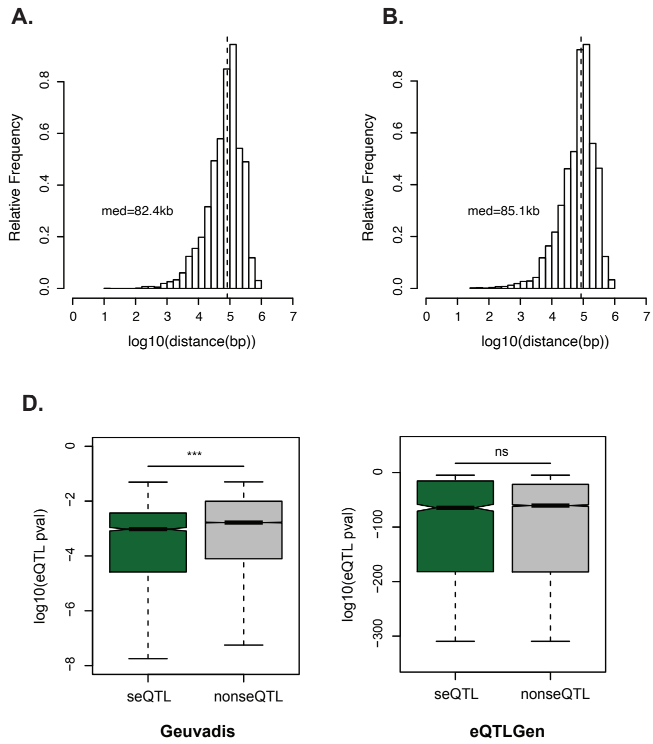


**Supplementary Figure 6. Impact of elncRNA splicing on *cis*-gene regulation in the human population.** Distribution of distance between genomic coordinate of seQTL and position of their associated elncRNA annotated transcriptional start sites (A) and enhancers (B). (C) Distribution of elncRNA sQTLs that are also associated with a *cis*-target expression (seQTL) as identified using a LCL population with 373 samples (Geuvadis) or blood samples (eQTLGen) with 31,684 samples. Distribution of elncRNA seQTLs and non-seQTLs (elncRNA sQTLs not associated with target expression in LCL population) associations as identified in LCL Geuvadis population (C) or in blood eQTLGen samples (D). Differences between groups were tested using a two-tailed Mann-Whitney *U* test. * $p < 0.05$; ** $p < 0.01$; *** $p < 0.001$; NS $p > 0.05$.

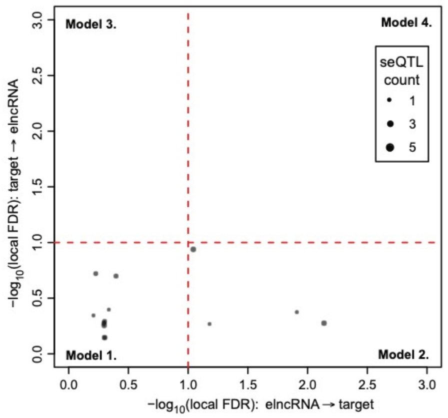


**Supplementary Figure 7. Impact of elncRNA splicing on chromatin signature associated with enhancer activity in the human population.** Scatterplot depicting the best joint elncRNA seQTLs (the most significant variant associated with elncRNA splicing) used to infer relationship between elncRNA splicing and *cis*-target expression levels using causal inference testing, as illustrated using local FDR associated with the four models (as illustrated in Figure 4B). Dotted black lines denote significance threshold at local FDR < 0.1.

#### SUPPLEMENTARY TABLE LEGENDS

Supplementary Table ST1. Genomic position of single- and multi-exonic LCL elncRNAs (hg19) and their respective number of exons.

Supplementary Table ST2. Single nucleotide polymorphisms (SNPs) at multi-exonic elncRNA splice sites and their respective targets, as well as SNPs jointly associated to both elncRNA and target expression levels. SNPs are denoted in the format snp_chr_position.

Supplementary Table ST3. Publicly available datasets used in the analysis.
